# The coumarin derivative X6632 is a pan-ID protein inhibitor that suppresses tumor growth by targeting cancer cells and the tumor-associated microvasculature

**DOI:** 10.64898/2026.08.05.742008

**Authors:** Lea S. Götz, Abhilash Deo, Sandra D. Scherer, Luigi D’Antonio, Hanna T. Weber, Georg Sedlmeier, Landy P. Torre Flores, Ulrike Kaiser, Elian Far, Ziv Raviv, Wilko Thiele, Sonja Thaler, Nicole Jung, Stefan Bräse, Caroline S. Hill, Alana L. Welm, Yuval Shaked, Boyan K. Garvalov, Jonathan P. Sleeman

## Abstract

Inhibitor of DNA binding (ID) proteins are key regulators of tumor cell stemness, therapy resistance and pathological angiogenesis in multiple cancer types and other diseases. Here, we characterize the coumarin-derived compound X6632 as a pan-ID inhibitor with dual activity against tumor cells and the tumor-associated microvasculature in a number of human and murine models. X6632 efficiently suppressed ID protein expression, inhibited the proliferation, migration, invasion of melanoma cells, and impaired multiple endothelial cell functions, including proliferation, migration, invasion, tube formation and sprouting in vitro. In back-to-back comparisons, X6632 exhibited an approximately ten-fold higher efficacy compared to the first-generation ID antagonist AGX51. In vivo, X6632 potently reduced pathological (neo)vascularization in established angiogenesis models, including oxygen-induced retinopathy and in Matrigel plug assays. It also significantly decreased blood vessel density in syngeneic melanoma models, delayed tumor growth and, when combined with immune checkpoint blockade, achieved superior tumor control compared with either monotherapy. Moreover, X6632 inhibited clonogenic growth in several breast cancer models, and robustly suppressed the growth of triple negative breast cancer in vivo, both in the highly aggressive 4T1 syngeneic model and in patient-derived xenografts. Collectively, these data establish X6632 as a second-generation, pan-ID protein inhibitor that can simultaneously target malignant cells and the tumor-supporting vasculature, and support the further pre-clinical development of the compound for the treatment of melanoma, breast cancer and potentially additional ID-dependent malignancies, as well as diseases driven by pathological neoangiogenesis.

## Introduction

Inhibitor of DNA binding proteins (ID proteins) are a family of class V helix-loop-helix (HLH) transcriptional regulators that play a multifaceted role in stem cell biology, angiogenesis, tumorigenesis, cancer cell stemness, and invasiveness [1–2]. ID proteins act as dominant-negative regulators of basic HLH transcription factors and form inactive heterodimers that cannot bind DNA, thereby suppressing cell differentiation and maintaining a proliferative cellular state [3–5]. Under physiological conditions, ID proteins are expressed in stem and progenitor cells, and are downregulated during cell differentiation. Accordingly, ID protein expression is tightly controlled by developmental cues, tumor suppressor pathways, post-transcriptional regulation [2], as well as growth factors such as TGFβ [6] and bone morphogenetic proteins (BMPs) [6]. In many types of cancer, including melanoma and breast cancer [7–13], inappropriate expression of ID proteins in tumor cells contributes to oncogenic processes, and can be driven by oncogenic signaling, constitutive growth factor stimulation, or compromised ubiquitin-proteasome pathway activity, each of which individually or collectively contribute to the pathological accumulation of ID proteins [2]. In addition to their cancer stemness promoting roles, ID proteins, and in particular ID1, have also been implicated in mediating resistance to chemotherapy, radiotherapy, and targeted agents [14–15].

ID proteins, especially ID1 and ID3, are also key pro-angiogenic regulators that help endothelial cells proliferate, migrate, survive, and form new blood vessels during development and in tumors [1]. They promote angiogenesis in part by repressing the expression of anti-angiogenic factors such as thrombospondin-1 and by amplifying VEGF-linked endothelial signaling pathways [16–17]. Genetic loss of ID1 and ID3 impairs vascular branching and vessel invasion, showing that these proteins are required for normal and pathological angiogenesis [18]. Cancer-associated aberrant neovascularization supports tumor growth, invasion, and metastasis, and represents a promising therapeutic avenue for anti-angiogenic and combination therapies [19–20]. Being expressed in both tumor and endothelial cells, ID proteins represent compelling candidates for targeted therapies [21].

Efforts to pharmacologically target ID proteins led to the identification of the small molecule AGX51, which promotes proteasomal degradation of ID proteins by disrupting their interaction with E proteins [22]. The compound reduced neovascularization in selected preclinical models [22] and suppressed breast tumor growth and metastasis in the lung when combined with paclitaxel, but not as a monotherapy [23], underlining the therapeutic relevance of ID proteins, while suggesting the need for more potent inhibitors of this protein family. Through a targeted chemical library screen, we previously reported the identification of a novel subclass of coumarin derivatives that target ID1 and ID3 [24]. The lead compound from this class, X6632, effectively suppressed extracellular matrix (ECM)-induced ID1 and ID3 expression in melanoma cell lines and significantly inhibited tumor initiation and growth in experimental animal models of melanoma.

In the present study, we show that X6632 is a pan-ID inhibitor that acts in both human and murine cells, and demonstrate that it has potent anti-angiogenic and anti-tumorigenic activity across multiple preclinical cellular and animal models. X6632 potently suppressed pathological (neo)vascularization in both tumor tissue and the retina, and exhibited approximately 10-fold higher efficacy than AGX51. In murine melanoma models, co-treatment with immune checkpoint inhibitors resulted in improved tumor control beyond that achieved with monotherapy. In addition, X6632 displayed pronounced efficacy in a range of breast cancer models, including the aggressive 4T1 syngeneic model as well as in patient-derived xenografts (PDX). These data position X6632 as a second generation pan-ID inhibitor and underline its potential for clinical application.

## Materials and Methods

### Cell lines and reagents

The murine melanoma cell lines B16F10 and RET, the human melanoma cell line A375, the murine breast cancer cell line 4T1 and the human breast cancer cell line SK-BR-3 were cultured in DMEM (Gibco, #11965092) containing 10% FCS (Gibco, #10270-106) and 1% penicillin/streptomycin (Gibco, #15140122). The murine melanoma cell line YUMM 1.7 was cultured in DMEM/F12 (Gibco, #11320033) containing 10% FCS, 1% penicillin/streptomycin and 1% MEM non-essential amino acid solution (NEAA) (Gibco, #11140035). Prior to in vivo transplantation of B16F10 and YUM1.7 cells for the PD-1 inhibition experiments, the cells were cultured in high glucose DMEM, supplemented with 10% FBS, 1% L-glutamine, 1% sodium-pyruvate and 1% penicillin–streptomycin. The human melanoma cell line HT144 was cultured in McCoy’s 5A medium (Gibco, #16600082) containing 10% FCS and 1% penicillin/streptomycin. The human breast cancer cell lines MCF7 and MDA-MB-231 were cultured in RPMI (Gibco, #11530586) with 10% FCS and 1% penicillin/streptomycin. The endocrine resistant human breast cancer cell line BT474 were cultivated in Advanced DMEM/F12 (Gibco, #11540006) supplemented with 10% FCS, 1% penicillin/streptomycin, 1% L-Glutamin (Gibco, #11500626) and 0.1% human insulin (Merck, #I9278). The human umbilical vein endothelial cells (HUVECs) (PromoCell GmbH, #C-12200) were cultured in Endothelial Cell Basal Medium 2 (PromoCell GmbH, # C-22111). Regular testing by Venor GeM kit (Minerva Biolabs) confirmed that the cells were mycoplasma-free. Compound X6632 was synthesized as previously published [24–25]. AGX51 (#HY-129241) was acquired from MedChemExpress. X6632 and AGX51 were dissolved in sterile DMSO (Carl Roth).

### Western blotting

Cells were lysed in 4% SDS, 125 mM TRIS-HCl (pH 6.8), sonicated, and boiled for 5 minutes. Proteins were separated on SDS-polyacrylamide gels and transferred to 0.2 µm nitrocellulose membranes (Amersham, Cytiva 10600002). The membranes were blocked with 5% nonfat milk and probed with rabbit monoclonal anti-ID1 (clone 195-14), rabbit monoclonal anti-ID2 (clone 9-2-8), rabbit monoclonal anti-ID3 (clone 17-3) antibodies (1:1000, ∼1µg/mL; BioCheck Inc), rabbit monoclonal anti-ID4 (clone 82-12) antibody (1:500, 2 µg/mL, BioCheck Inc) and anti-Hsp90 (clone C45G5) antibody (1:10000, 1.3ng/mL) (Cell Signaling Technology, 4877T), anti-Vinculin (clone hVIN-1) antibody (1:200000, 0,046 µg/mL) (Merck, #V9131) or anti-β-actin mouse monoclonal (clone AC-15) antibody (1:10000, 0.21 µg/mL; Sigma Aldrich) as loading controls. After washing and incubation with horseradish peroxidase (HRP)-conjugated secondary antibodies: polyclonal goat anti-rabbit (P044801-2, Dako) and polyclonal goat anti-mouse (P044701-2, Dako). Antibody-bound proteins on the membranes were detected using enhanced chemiluminescence (ECL) (Thermo Fisher Scientific). Brightness and contrast of the images were adjusted using GIMP software.

### Proliferation assay

For proliferation assays cells were seeded in 50 µL of suitable medium into the wells of 96-well plates (4.5 x 10^3^ for HUVEC and 1 x 10^3^ for melanoma cell lines). The next day, the cells were stimulated for 48h using different concentrations of X6632 or AGX51 ranging from 0.15 µM to 160 µM. Cell numbers were quantified using the CyQuant assay (Thermo Fisher Scientific, #C35006) according to the manufacturer’s instruction with a TECAN Infinite M Nano^+^ microplate reader. A total of four replicates were performed for each condition. The IC_50_ was calculated using GraphPad Prism.

### Cell exclusion zone migration assay

For cell-exclusion migration assays, 5 x 10^5^ HUVECs in 70 µL medium were seeded into each well of a 2-well silicone culture inserts (ibidi GmbH, #80209). The next day, the inserts were removed carefully and 2 mL of medium containing either 0.1% DMSO or suitable concentrations of X6632 or AGX51 (2.5 µM) was added to the wells. The same area was imaged using a Leica DM600 microscope immediately after insert removal (T_0_) and 16 hours after removal (T_16_). A total of six replicates were performed for each condition. Images were analyzed using ImageJ/Fiji [26].

### Transwell migration and invasion assays

Transwell cell culture inserts (Corning, #3422) with a pore size of 8 µm were used for Boyden chamber migration and invasion assays. Migration assays were performed by seeding 1 x 10^4^ cells in 100 µL cell line-specific medium supplemented with 1% FCS on top of the membrane. After an 8-hour incubation to allow cell attachment, the cell line-specific medium in the lower chamber was exchanged for medium supplemented with 10% FCS to induce a chemoattractant gradient and stimulate migration, in the presence of 0.1% DMSO or suitable concentrations of X6632 or AGX51 (2.5 µM for HUVEC, 5 µM for melanoma cell lines). Simultaneously, a further 100 µL of cell line-specific medium supplemented with 1% FCS and 0.2% DMSO or twice the desired final concentration of the stimulants (for HUVEC 5 µM, for melanoma cell lines 10 µM) were added to the upper compartment. After 24 hours, cells were fixed in 4% PFA and stained with DAPI solution (5 µg/mL in PBS). For invasion assays, the membrane was coated with high concentration, phenol red-free Matrigel (Corning, #354263) at a concentration of 0.6 mg/mL in PBS), and incubated at 37°C overnight. Cell seeding and stimulation were performed as described for migration assays, with invasion times adjusted for each cell type as indicated in the figure legends.

Total cells were imaged using a Leica DM600 microscope. Cells in the upper compartment were then removed with cotton swabs and the invaded cells on underside of the membrane were imaged on a Leica DM600 microscope. Images were analyzed using ImageJ/Fiji.

### Tube formation assay

For tube formation assays, 300 µL of high concentration, phenol red-free Matrigel (Corning, #354263) diluted in PBS to a concentration of 10 mg/mL was added to the wells of a 24-well plate and incubated at 37°C for 30 minutes to allow the Matrigel to polymerize. Subsequently, 6 x 10^4^ HUVECs (P3 to P8) in 0.3 mL medium containing either 0.1% DMSO or 5 µM X6632 or AGX51 were added onto the solidified Matrigel. After 6 hours the tubular network was imaged on a Leica DM600 microscope at 5x magnification. Quantification of total segment length, number of master junctions, number of meshes and total mesh area was performed using ImageJ/Fiji and the Angiogenesis Analyzer plugin [27].

### Spheroid sprouting assay

A methylcellulose-based hanging drop method was used for spheroid formation using HUVECs. A 1.2% (w/v) methylcellulose solution was prepared in DMEM/F12 Advanced medium (Gibco, #12634010). The methylcellulose solution was mixed with HUVEC growth medium at a ratio of 9:1 (v/v), then 1.6 × 10^5^ cells were suspended in the mixture. For hanging drop culture, 25 μL of the cell suspension was seeded onto inverted 15 cm dish lids and incubated for 24 hours. The next day, spheroids were collected in 1.5 mL growth medium and 1 mL methylcellulose solution and mixed 1:1 (v/v) with a neutralized rat collagen solution (3.2 mg/mL rat collagen (Serva, #47256.01), 1x HBSS and 2.3 mg/mL NaHCO_3_ (pH 7), adjusted with NaOH using phenol red as an indicator). The collagen-embedded spheroids were seeded into pre-warmed 24-well plates and incubated at 37°C. After 30 minutes, 200 µL of medium containing 25 ng/mL VEGF-A isoform 164 (Reliatech, M30-004) and either 0.1% DMSO or 10 µM of X6632 or AGX51 was added to the wells. The spheroids were imaged after 24 hours using a Leica DM600 microscope (20x magnification), and the number of sprouts and sprout length were analyzed using ImageJ/Fiji.

### Clonogenic assay

For colony formation assays, MCF7, BT474, MDA-MB-231 and SK-BR-3 cells were seeded at a density of 1-7.5 x 10^3^ cells/well in 6-well plates, then treated with increasing concentrations of X6632 for 14-21 days. The medium containing the compound was replaced on a weekly basis. At the end of the experiment, cells were fixed with 20% (v/v) methanol and the colony formation was visualized after staining with 0.5% crystal violet (w/v) in 20% methanol (v/v). Plates were photographed and colony formation was quantified using ImageJ/Fiji as area coverage, with normalization against DMSO-treated controls.

### Experimental mice

Animal experiments were approved by the local regulatory authorities (license numbers: AZ 35-9185.81/G-296/20 and AZ 35-9185.81/G-98/21), and were performed according to local legal requirements. BALB/c and C57/Bl6RJ mice were purchased from Janvier. Mice were kept in groups of four in type III and type IIL polysulfon IVC cages (Tecniplast, Hohenpeißenberg, Germany) containing soft bedding (Mini Chips LTE E-003 L-10 ABEDD B, Ssniff GmbH, Soest, Germany). Mice were also provided with nesting material (Pura Cocoons Soft Cotton Cylinder, Code 211–1240 HV/MG/0822, Labodia, Switzerland), a mouse house and tubes (Play Tunnels: CS3B01213–1041) made of cardboard and wooden sticks (Nagehölzer, H0234-NGS, Ssniff and Pura Sticks, Code 213–1001, Labodia, Switzerland) for gnawing. The cages were changed exclusively under laminar flow conditions. Rat/mouse maintenance complete feed (Ssniff) and tap water were provided ad libitum. The specific pathogen-free housing was kept at 20–22°C and 45–60% humidity on a 06:00–18:00 h light cycle. Routine health monitoring was carried out in accordance with FELASA recommendations [28]. Parasitological tests were performed in the laboratory of the Core Facility Preclinical Models, bacteriological tests in the Institute of Microbiology, Klinikum Mannheim and serological and PCR tests at BioDoc, Labor für Biomedizinische Diagnostik, Hannover and GIM (Gesellschaft für innovative Mikroökologie mbH, Michendorf, Germany).

### Oxygen-induced retinopathy mouse model

One-week-old C75BL/6JRj mice (P7) were kept in hyperoxia chambers (75% oxygen) for 5 days (P12). On P12, mice received a single intravitreal injection in each eye of X6632 (injection volume: 1 µL, final concentration: 0.6 mg/kg), or injections with the control substances DMSO and PBS (injection volume: 1 µL). At P17, mice were sacrificed and the eyes were snap frozen in liquid nitrogen. Eyes were fixed in 3.7% formalin overnight at 4°C and processed the next morning. Retinas were dissected, permeabilized in ice-cold methanol, and incubated in blocking buffer (1% (w/v) BSA (Carl Roth), 5% (v/v) normal serum (Gibco, #C06SB) in PBS/Triton X-100 (0.5%) (Carl Roth)) overnight at 4°C. After washing in PBS/Triton X-100 (0.5%), the retinas were incubated in isolectin B4 (1:100; Thermo Fisher Scientific, #I21411) for 72 hours at 4°C, and DAPI (5 µg/mL) was added for the last night. Retinas were mounted with Fluoromount G (Thermo Fisher Scientific, 00-4958-02), and were imaged using a Zeiss LSM800 confocal microscope. Analysis was performed for each quadrant using ilastik [29] and ImageJ/Fiji.

### Matrigel plug assay

A volume of 500 µL of growth factor-reduced, phenol red-free Matrigel (Corning, #356231) (concentration: 7 mg/mL) was mixed with 250 ng/mL VEGF-A isoform 164 (Reliatech, M30-004) and injected subcutaneously into the flank of eight-week-old BALB/cJRj mice. After the Matrigel injection, the mice received intraperitoneal injections of X6632 (18 mg/kg) or the corresponding volume of DMSO for 7 days, and were sacrificed on the day after the last injection. Plugs were cut in half for paraffin- and cryosectioning. Cryosections 20 µm thick were used for immunohistochemical staining. Sections were fixed in ice-cold methanol/acetone (1:1), treated with 0.3% H_2_O_2_ (Carl Roth) for a peroxidase block, and then blocked using 1% BSA (Carl Roth) and 10% normal serum (Gibco, #16210-072) in PBS. Sections were stained for blood vessels overnight at 4°C with the anti-CD31 antibody clone MEC13.3 (BD Biosciences, #550274) diluted 1:250 (62.5 ng/mL). Biotinylated secondary antibodies (Vector Laboratories) were applied for 45 minutes and the signal was developed using ABC reagent (Biozol Diagnostics, #ZE0906) and NovaRed substrate (Vector laboratories, #sk4800) according to the manufacturer’s instruction. Hematoxylin counterstaining was performed and sections were embedded in Eukitt. Four sections per plug were imaged using a Zeiss light microscope AXIO Imager Z1. Analysis was performed with ImageJ/Fiji.

### Tumor cell injection

B16F10, RET, YUMM 1.7 and 4T1 tumor cells (2 x 10^6^ in 100µL PBS) were injected subcutaneously into the flank of syngeneic mice. Mice were regularly monitored for body weight and tumor growth. Daily treatment with X6632 (18mg/kg) or an equivalent volume of DMSO i.p. was started as soon as a tumor was detectable (∼2-5 mm in one dimension). The day after the last injection, the mice were sacrificed. The tumor with skin was cut in half for paraffin- and cryosectioning. The ipsi- and contralateral axillary and inguinal lymph nodes were collected, and snap frozen in liquid nitrogen. Lungs were harvested and fixed in 3.7% formalin. Cryosections 12 µm thick were used for immunofluorescence staining. Sections were fixed in ice-cold methanol/acetone (1:1) and blocked in blocking buffer (1% (w/v) BSA, 10% (v/v) normal serum (Gibco, #C06SB) in PBS)). Sections were stained overnight at 4°C with the anti-CD31 antibody clone MEC13.3 (BD Biosciences, 550274) diluted 1:250 (62.5 ng/mL), followed by incubation for 1 hour with fluorescently-labelled secondary antibody (Thermo Fisher Scientific, #A-110081) diluted 1:1000 (Thermo Fisher Scientific) and DAPI (5 µg/mL), then mounting in FluoromountG (Thermo Fisher Scientific, #00-4958-02). Four sections per tumor were imaged using a Zeiss Light Microscope AXIO Imager D1. Analysis was performed with ImageJ/Fiji.

### Experimental lung metastasis model

The tail vein injection experiments were performed at the Francis Crick Institute under the Animals (Scientific Procedures) Act 1986 in accordance with UK Home Office license (Project License PP5869518) and the EU Directive 2010. The Home Office licenses underwent full ethical review and approval by the Francis Crick Institute’s Animal Ethics Committee. The mice were housed in constant temperature, humidity, and pathogen-free controlled environment (25 °C ± 2 °C, 50–60%) cages with a standard 12 h light / 12 h dark cycle and were allowed access to food and water ad libitum. 4T1 breast cancer cells from the Cell Science Facility at the Francis Crick Institute, certified mycoplasma negative, were resuspended in sterile PBS and injected intravenously into immunocompetent BALB/c recipient mice via the tail vein. Each mouse received 5 x 10^5^ cells in a final volume of 0.1 mL. A total of 16 mice were divided into two groups: a control group receiving DMSO and a treatment group receiving the ID1/ID3 inhibitor X6632. X6632 was administered 5 days a week for 10 days by intraperitoneal injection at a dose of 18 mg/kg. Ten days after tumour cell injection, mice were culled by cervical dislocation, death was confirmed by post-mortem examination, and lungs were collected and fixed in formalin for downstream analysis of metastatic burden by quantification of metastatic nodules in HE-stained lung sections.

### Combination treatment of X6632 and anti-PD-1

B16F10 and YUMM 1.7 tumor cells (1 x 10^6^ in 100 µL PBS) were injected subcutaneously into the flank of syngeneic mice. Mice were regularly monitored for body weight and tumor growth. Daily treatment with X6632 (18 mg/kg) or an equivalent volume of DMSO was started when tumors reached a detectable size (∼30-50 mm^3^). In addition, intraperitoneal injections with isotype control rat IgG (BioXCell or IchorBio) or anti-PD1 antibody (RPM1-14 clone, BioXCell or IchorBio) at 100 µg in 100 µL PBS were performed every third day, starting from the first day of treatment with DMSO or X6632. Flow cytometry analysis of the tumor tissue was performed as previously described (Benguigui et al., 2024).

### PDX HCI-043 Model

HCI-043 tumor fragments were thawed and implanted into the inguinal mammary fat pad of 3-4 weeks old female immune-compromised NRG mice (NOD.Cg-*Rag1^tm1Mom^ Il2rg^tm1Wjl^*/SzJ, Jackson Laboratory cat # 007799). Mice were monitored for body weight and tumor growth twice a week. Mice were enrolled into the study when tumor size reached ∼100 mm^3^, and randomly assigned to treatment and control group. Daily i.p. treatment with X6632 (18 mg/kg) or DMSO was started as soon as mice were enrolled, and mice received 24 daily injections. The day after the last injection, the mice were sacrificed. Tumors were harvested, cut in half, and snap frozen for cryo blocks, or fixed in 4% PFA, processed and embedded for histology blocks. Cryosections 12 µm thick were used for immunofluorescence staining. Sections were fixed in ice-cold methanol/acetone (50/50) and blocked in blocking buffer (1% (w/v) BSA, 10% (v/v) normal serum (Gibco, #C06SB) in PBS)). Sections were stained with an anti-CD31 antibody (clone MEC13.3) and imaged as described above.

### Flow cytometry acquisition and analysis

Flow cytometry analysis of the tumor tissue was performed as previously described [30]. Flow cytometry stainings were performed using surface markers to identify different immune cell types as indicated in Table S1. All antibodies were purchased from BD Biosciences, USA or BioLegend, USA: APC/Cyanine7 rat anti-mouse CD45 antibody (Clone 30-F11), Cat# 557659; BUV395 rat anti-mouse CD45R/B220 antibody (Clone RA3-6B2), Cat# 563793; RB705 rat anti-mouse CD4 antibody (Clone GK1.5), Cat# 570257; BUV496 rat anti-mouse CD8a antibody (Clone 53-6.7), Cat# 569181; APC rat anti-mouse CD25 (Clone PC61), Cat# 557192; RB613 Hamster Anti-Mouse CD11c antibody (Clone N418), Cat# 758187; RB545 rat anti-CD11b antibody (Clone M1/70), Cat# 569255; PE/Cyanine7 anti-mouse CD335 (NKp46) antibody (Clone 29A1.4), Cat#137617; BV711 hamster anti-mouse CD49b (Clone HMα2), Cat#740704; B744 rat anti-mouse Ly-6C (Clone AL-21), Cat# 570492; V450 rat anti-Mouse Ly-6G (Clone 1A8), Cat# 560603; RB780 rat anti-mouse F4/80 (Clone T45-2342), Cat# 569223. The antibodies were used in accordance with the manufacturer’s instructions. The samples were processed using a FACSymphony A5 SE flow cytometer and the data were analyzed with FlowJo V.10 software (FlowJo, Ashland, Oregon, USA).

### Statistical analysis

Statistical analyses were performed using GraphPad Prism. Data are presented as mean ± standard error of the mean (SEM). For comparing two groups, a two-tailed Student’s t-test was used. For comparing more than two groups, one-way ANOVA with Dunnett’s multiple comparisons tests was used. Tumor growth curves were compared using two-way repeated measures ANOVA with Tukey’s multiple comparisons tests. Statistical significance was defined as follows: p < 0.05 (*), p < 0.01 (**), p < 0.001 (***), p < 0.001 (****); non-significant differences are indicated as “ns” (p > 0.05).

## Results

### X6632 has strong inhibitory effects on the proliferation, migration and invasion of human and mouse melanoma cells

Building on our previous findings [24], we undertook a detailed characterization of the effects of X6632 on the behavior of a murine and human melanoma cells. First, we tested increasing concentrations of X6632 on the ID1 and ID3 protein levels of B16F10 murine melanoma and A375 human melanoma cells. A concentration of 7.5 µM X6632 potently reduced ID1 and ID3 protein levels in both cell lines (Figure 1A and E). Next, we tested the effects of X6632 and AGX51 on melanoma cell proliferation using an in vitro CyQuant assay (Figure 1B and F). The IC_50_ values for X6632 were approximately 3 µM for both cell lines, demonstrating a potent inhibitory activity of the compound. By contrast, AGX51 exhibited significantly higher IC_50_ values of 46.17 µM for B16F10 and 12.66 µM for A375. In addition, we conducted transwell migration and invasion assays with both melanoma cell lines using 5 µM X6632 and AGX51, or DMSO as a control. X6632 strongly and significantly inhibited migration and invasion in B16F10 and A375 cells (Figure 1C, D, G, H). By contrast, AGX51 had no effect on the migration and invasion of A375 cells (Figure 1G, H), had no effect on the migration of B16F10 cells (Figure 1C), and led to only a partial decrease in the invasion of B16F10 cells (Figure 1D). Taken together, the in vitro data demonstrated a consistent and potent ID protein inhibition and anti-tumor effects of X6632 in both human and murine melanoma cells.

**Figure 1:**
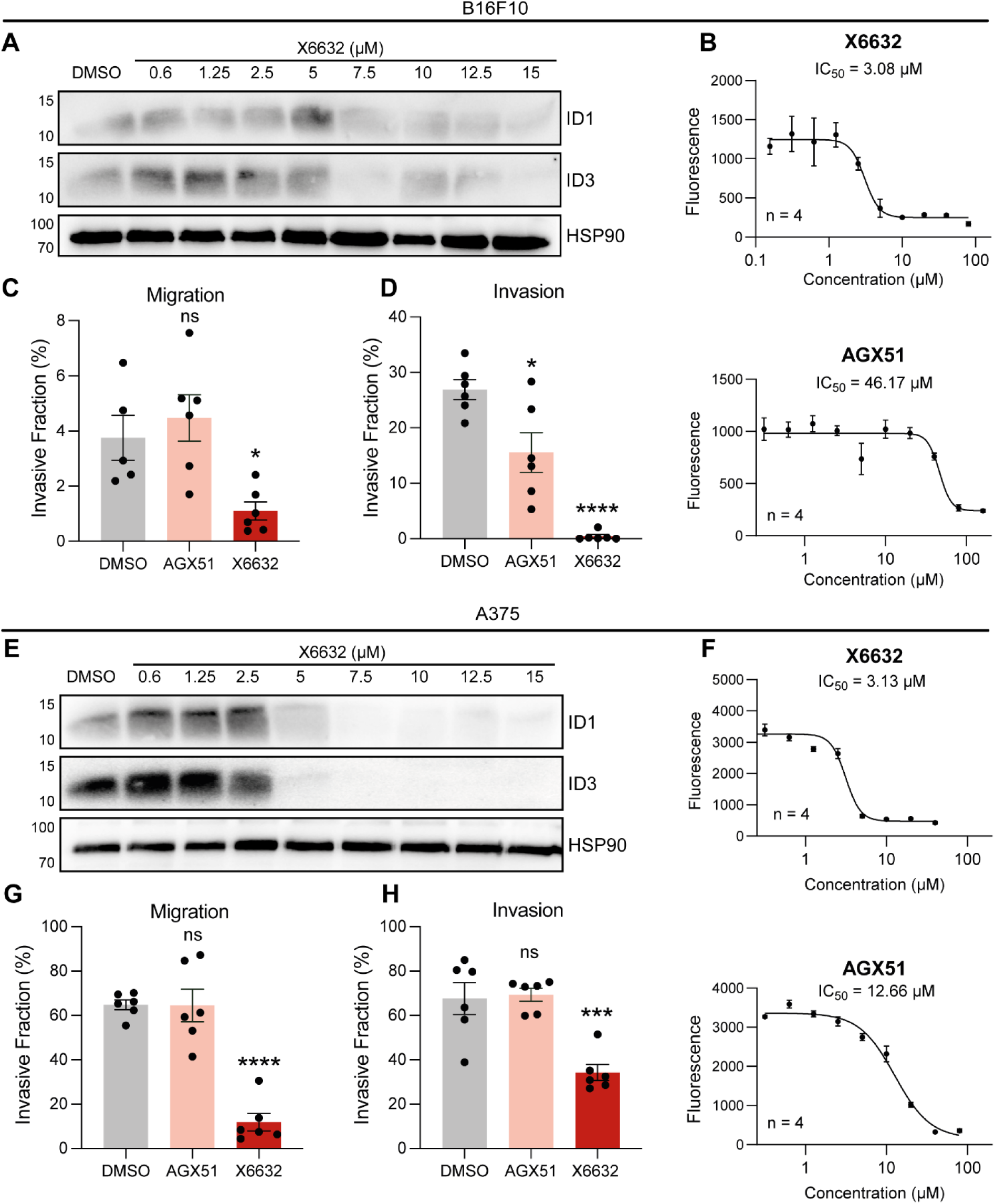
Potent anti-cancer activity of X6632 in human and murine cells in vitro. **A** Western blot-based analysis of dose-dependent effects of X6632 on ID1/ID3 levels in murine melanoma cell line B16F10. **B** Proliferation assay with the murine melanoma cell line B16F10 treated with X6632 or AGX51 for 48 hours (n = 4). **C** Transwell-based migration assay, using uncoated inserts with B16F10 cells treated with 5 µM X6632 or AGX51 for 48 hours (n = 6). **D** Transwell-based invasion assay using Matrigel coated inserts with B16F10 cells treated with 5 µM X6632 or AGX51 for 96 hours (n = 6). **E** Western blot-based analysis of dose-dependent effects of X6632 on ID1/ID3 levels in human melanoma cell line A375. **F** Proliferation assay with the human melanoma cell line A375 treated with X6632 or AGX51 for 48 hours (n = 4). **G** Transwell-based migration assay, using uncoated inserts with A375 cells treated with 5 µM X6632 or AGX51 for 20 hours (n = 6). **H** Transwell-based invasion assay using Matrigel coated inserts with B16F10 cells treated with 5 µM X6632 or AGX51 for 24 hours (n = 6). *P ≤ 0.05; ***P ≤ 0.001; ****P ≤ 0.0001

### X6632 has potent anti-angiogenic effects in multiple in vitro assays

Given the role of ID proteins in angiogenesis, we next wanted to investigate the anti-angiogenic potential of X6632. To this end, we first examined the effect of the compound on the proliferation, migration, invasion, tube formation and sprouting of human umbilical vein endothelial cells (HUVECs) and compared it to the pan-ID antagonist AGX51 [22]. W determined the dose-dependent effects of the two compounds on the protein levels of ID1 and ID3 in HUVECs using Western blotting. A concentration of 0.3 µM X6632 was sufficient to potently reduce the ID1 and ID3 protein levels within 24 hours (Figure 2A). By contrast, 20 µM AGX51 was required to downregulate ID1 and ID3 protein levels. In addition, the effects of X6632 on cell proliferation were compared with those of the pan ID-antagonist AGX51 in an in vitro proliferation assay (CyQuant, Figure 2B), revealing that X6632 had an IC_50_ value that was more than 10-fold lower than that of AGX51 (0.91 µM vs 9.86 µM).

**Figure 2:**
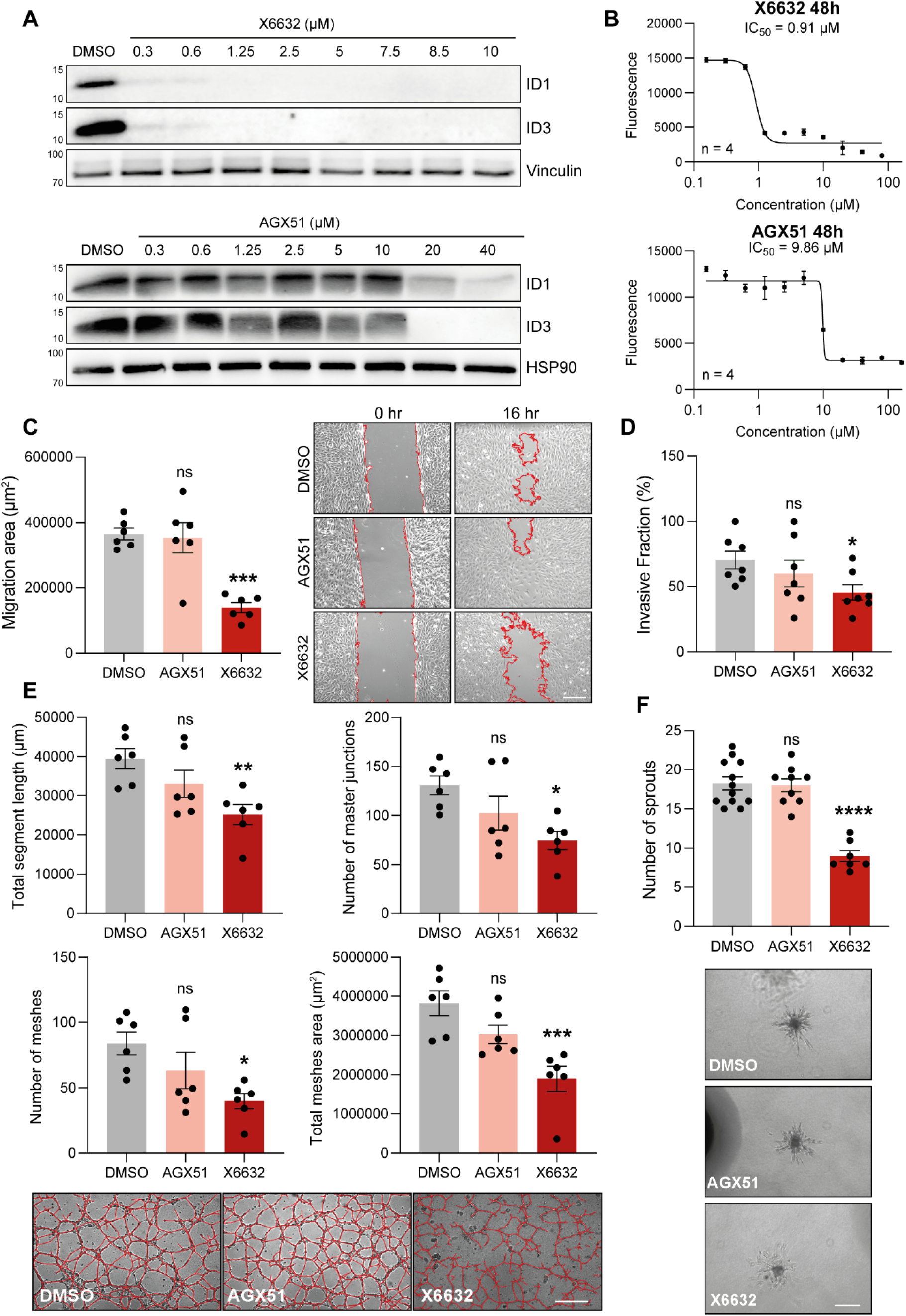
Potent anti-angiogenic activity of X6632 in multiple in vitro assays. **A** Western blot-based analysis of dose-dependent effects of X6632 on ID1/ID3 levels in HUVECs. **B** Viability assay after X6632 and AGX51 treatment of HUVECs for 72 hours (n = 4). **C** Silicone insert-based assay for HUVEC migration for 16 hours with 2.5 µM X6632 or AGX51 (n = 6) Representative images are shown on the right, with the cell-free areas encircled in red (scale bar = 200 µm). **D** Transwell-based Invasion assay with HUVECs for 72 hours with 2.5 µM X6632 or AGX51 (n = 6). **E** Quantification of total segment length, number of master junctions, number of meshes and total mesh area in a tube formation assay with HUVECs plated on Matrigel and treated for 6 hours with 5 µM X6632 or AGX51 (n = 6). Representative images from each condition are shown at the bottom, with the tubular structures traced in red (scale bar = 200 µm). **F** Quantification of the number of sprouts in a spheroid-based sprouting assay with HUVECs treated for 24 hours 10 µM X6632 or AGX51 (n = 12 for DMSO, n = 9 for AGX51, n = 7 for X6632). Representative images from each condition are shown at the bottom (scale bar = 200 µm). ns, not significant (P > 0.05); *P ≤ 0.05; **P ≤ 0.01; ***P ≤ 0.001; ****P ≤ 0.0001.

We next assessed the impact of X6632 on angiogenesis-related properties of HUVECs in several functional assays. First, the effects of X6632 and AGX51 on cell migration were tested using a silicone insert-based migration assay with DMSO or 2.5 µM of X6632 or AGX51 for 16 hours. While X6632 significantly reduced HUVEC migration within 16 hours compared to the DMSO control, AGX51 had no inhibitory effect (Figure 2C). The influence of X6632 and AGX51 on endothelial cell invasion was then assessed using Matrigel-coated transwell inserts. Exposure of the HUVECs to 2.5 µM X6632 for 72 hours significantly diminished cell invasion compared to control conditions with DMSO. By contrast, no suppressive effect was observed with 2.5 µM AGX51 (Figure 2D).

To further examine the anti-angiogenic potential of X6632 and AGX51, we used a tube formation assay, in which HUVECs seeded on Matrigel form capillary-like structures. The freshly seeded cells were exposed to DMSO or 5 µM of X6632 or AGX51 for 6 hours, then a quantitative analysis of parameters that provide information on network complexity was performed, including the total segment length, the number of master segments, as well as the number of meshes and nodes. X6632 significantly attenuated all parameters, whereas AGX51 did not elicit significant changes, with only a slight trend towards reduction compared to the DMSO control (Figure 2E). Finally, we tested the effects of X6632 compared to AGX51 on endothelial cell sprouting, using a spheroid-based assay [31] (Figure 2F). HUVEC spheroids were cultured with VEGF-A (25 ng/mL) to induce sprouting, and were treated with either X6632 or AGX51 (10 µM), or DMSO as a solvent control. After 24 hours of stimulation, the number of sprouts per spheroid was significantly reduced with X6632, while no effect was observed with AGX51. In summary, X6632 demonstrated significantly greater efficacy than AGX51 in suppressing the angiogenic properties of endothelial cells in vitro.

### X6632 is a pan-ID inhibitor

X6632 was identified in a chemical library screen for compounds that inhibit ID1 and ID3 expression [24]. To determine whether X6632 is a pan-ID inhibitor, similar to AGX51, we investigated the impact of X6632 on the expression of all four ID proteins in two human melanoma cell lines and HUVECs. X6632 potently reduced expression of all four ID proteins at a concentration of 10 μM over a 24-hour period (Figure 3). These data demonstrate that X6632 is a pan-ID inhibitor rather than a selective ID1/ID3 antagonist.

**Figure 3:**
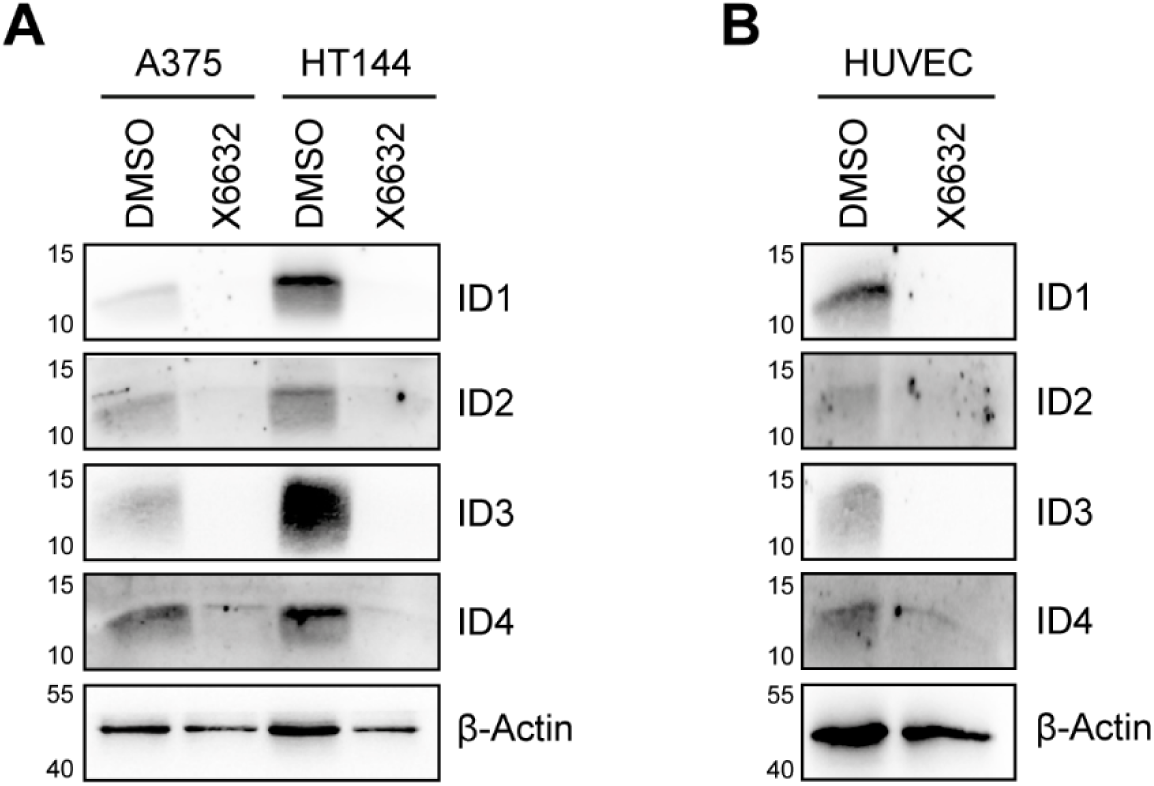
X6632 inhibits the expression of all four ID proteins. A375 and HT144 human melanoma cells (**A**) or HUVECs (**B**) were treated with DMSO or 10 μM X6632 for 24 hours. HT144 cells were additionally stimulated with BMP4 (20 ng/mL) to induce ID protein expression. The expression of the four ID proteins was determined using Western blotting.

### X6632 inhibits angiogenesis in vivo

Our in vitro data with HUVECs suggests that X6632 inhibits the angiogenic properties of endothelial cells. To determine whether this translates into an effect on angiogenesis in vivo, we employed three independent angiogenesis models. First, we used a mouse model of oxygen-induced retinopathy (OIR) to demonstrate the efficacy of X6632 as an angiogenesis inhibitor in vivo (Figure 4A). The role of ID proteins in ocular pathological neovascularization in the OIR model has been demonstrated genetically by using ID1 or ID3 knockout mice [22]. Newborn mice have an immature retinal vasculature that develops postnatally [32–33]. In the OIR mouse model, neonatal mice (P7) are exposed to hyperoxia (75% oxygen) for 5 days (P7 to P12), which induces degeneration and regression of retinal vessels in the central zone, called vaso-obliteration (VO). Upon return to ambient oxygen at P12, the relative hypoxia in the retina leads to overexpression of angiogenic growth factors such as VEGF-A, causing sprouting from capillaries in the retinal periphery that typically result in pathological neovascularization, in the form of neovascular tufts that grow towards the vitreous body [34] (Figure 4B).

**Figure 4:**
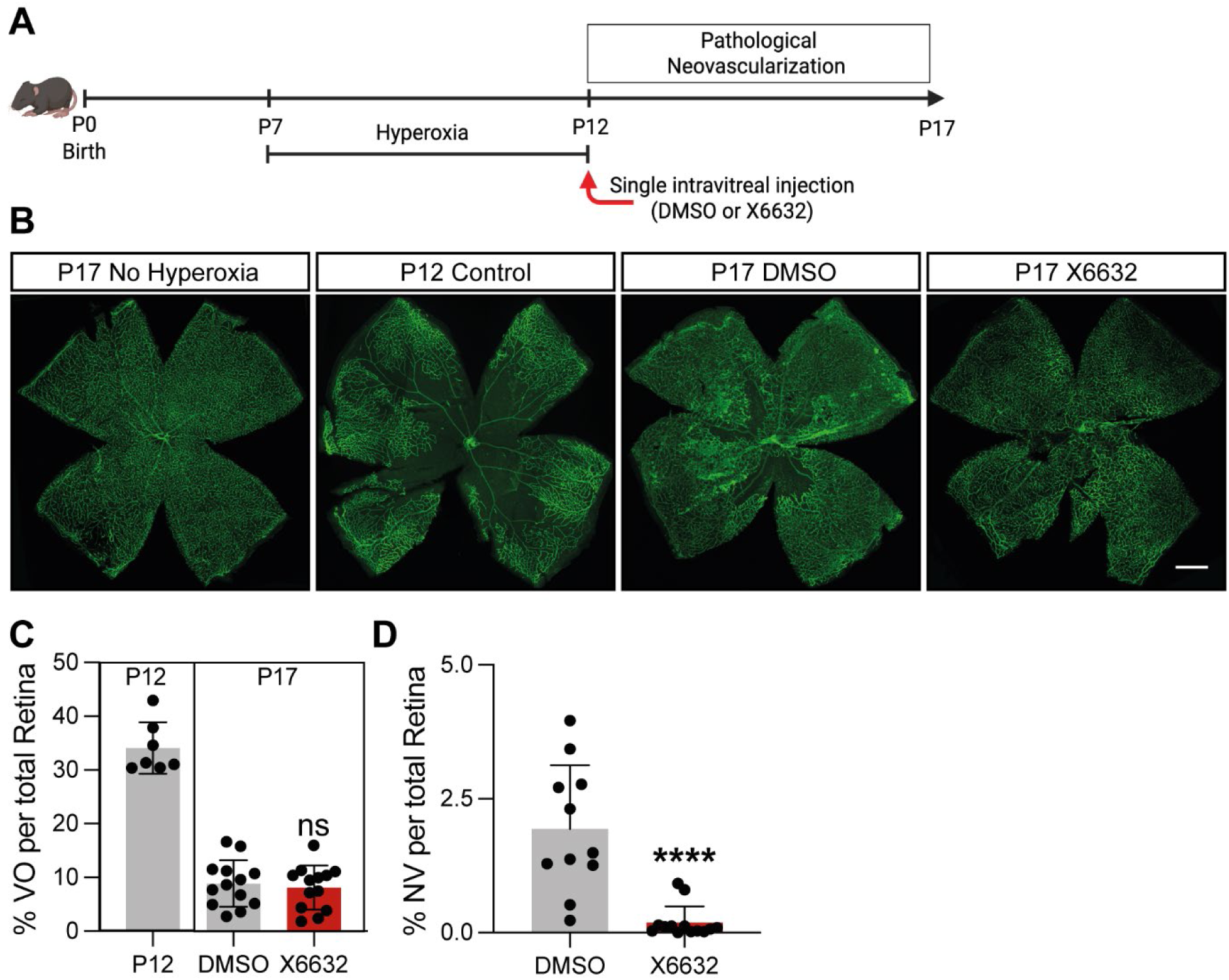
X6632 selectively protects against pathological neovascularization in oxygen-induced retinopathy. **A** Schematic of the OIR model: Hyperoxia (75% O_2_, P7-P12) induces central retinal vessel loss; return to normoxia (20% O_2_) triggers VEGF-A-driven pathological neovascular tuft formation. Single intravitreal injection of DMSO or X6632 at P12 (injection volume: 1 µL, final concentration: 0.6 mg/kg). Endpoint: P17. **B** Representative images of retinal vessel staining with isolectin B4 of P17 control retina (not subjected to hyperoxia), P12 retina after hyperoxia and P17 retinas after hyperoxia and treatment with DMSO or X6632 (scale bar = 500 µm). **C** Quantification of the vaso-oblitaration (VO) area in P12 animals after hyperoxia, and in The DMSO and X6632 treated groups at P17 (n = 7 for P12 control; n = 13 for DMSO and X6632). **D** Quantification of the relative area with pathological neovascularization (NV) at P17 after treatment with DMSO or X6632 (n = 11 for DMSO, n = 13 for X6632). ns, not significant (P > 0.05); ****P ≤ 0.0001.

Newborn mice exposed to hyperoxia from P7 to P12 were returned to ambient oxygen and given an intravitreal injection of X6632 or DMSO as a control. The impact on VO and pathological neovascularization was then assessed at P17. The VO zone decreased in both experimental groups at P17 compared to P12 (Figure 4C), indicating that revascularization of the retina was not affected by X6632 treatment. However, X6632 strongly and significantly reduced the amount of pathological neovascularization at P17 compared to the vehicle control (Figure 4D). These findings demonstrate that X6632 protects the retina from pathological neovascularization, without interfering with physiological retinal revascularization.

Next, we determined whether X6632 is able to suppress growth factor-induced angiogenesis in vivo using the Matrigel plug assay [35]. Growth factor-reduced Matrigel was mixed with VEGF-A (250 ng/mL) to induce blood vessel ingrowth into the plug, and was then injected subcutaneously into the flank of mice. Animals were treated daily for 7 days with either DMSO or X6632 (18 mg/kg, ip) (Figure 5A), and the blood vessel density in the plug and the adjacent cutaneous tissue was quantified the day after the last injection using immunohistochemistry. This analysis revealed that X6632 treatment significantly reduced the blood vessel density in the plug and in the adjacent cutaneous tissue compared to the vehicle treatment (Figure 5B, C).

**Figure 5:**
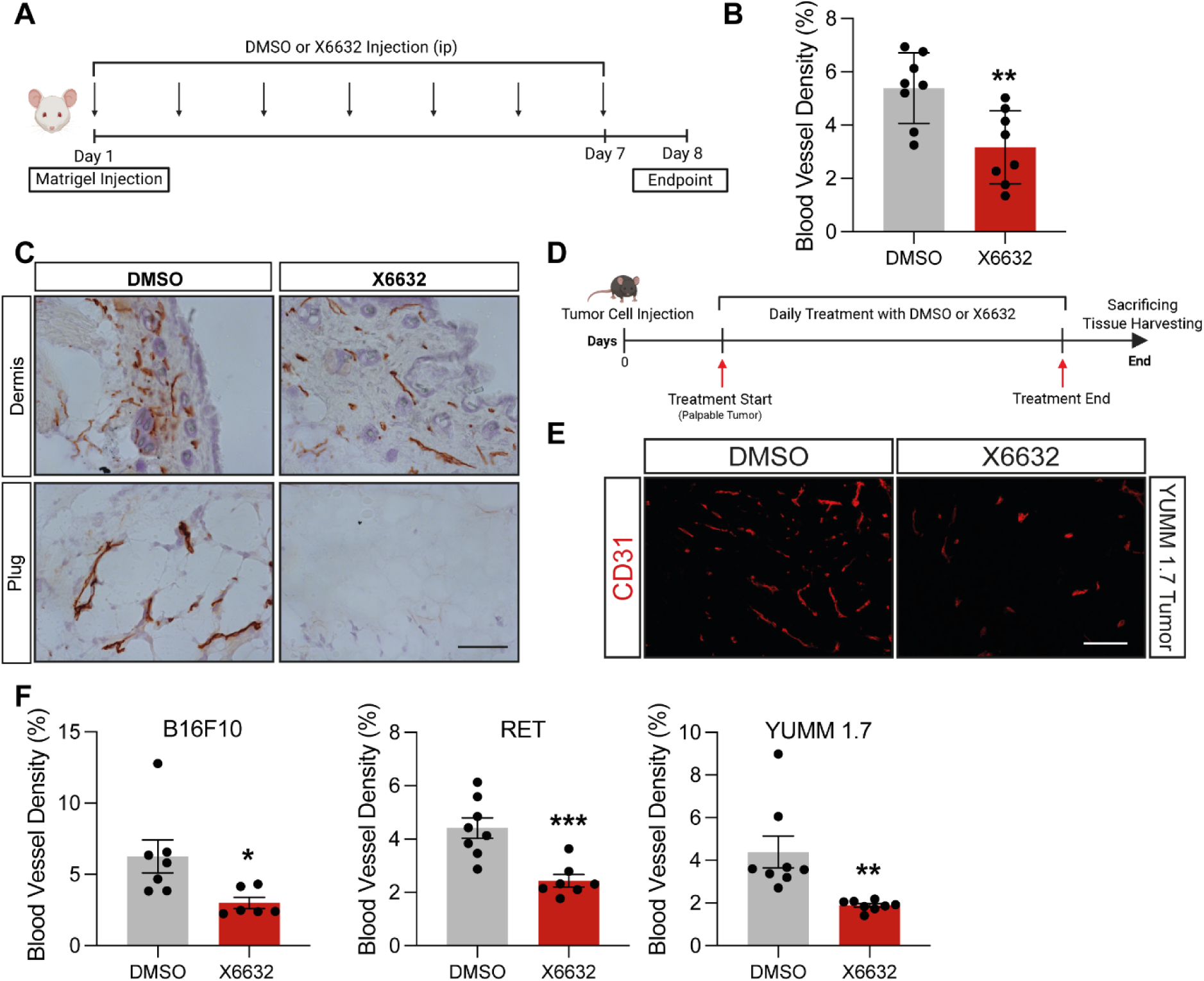
Anti-angiogenic properties of X6632 in preclinical animal models. **A** Schematic of Matrigel plug assay: growth-factor-reduced Matrigel mixed with VEGF-A (250 ng/ml) was subcutaneously injected into BALB/c mice and treatment on 7 consecutive days with X6632 or DMSO was performed. 8 days after injection the plugs were excised and formation of blood vessels into the Matrigel was analyzed. **B** Representative images of IHC staining on cryosections with an anti-CD31 antibody and a hematoxylin counterstaining (scale bar = 100 µm) **C** The blood vessel density in the skin as well as in the plug was analyzed (n = 8). **D** Schematic of tumor models: 2 x 10^6^ cells tumor cells were injected subcutaneously into the flank of mice. Daily treatment with DMSO or X6632 (ip, 18mg/kg) started once the tumors reached a size of 50 mm^3^. Thereafter B16F10 tumor-bearing mice received 9 daily injections of X6632, while RET and YUMM 1.7 tumor-bearing mice received 11 injections. **E** Representative images of blood vessel density (anti-CD31) in tumor cryosections of YUMM 1.7 tumors of animals treated with DMSO or X6632 (scale bar = 100 µm). **F** Quantification of blood vessel density in the tumors after treatment with DMSO or X6632. (B16F10: n = 7 DMSO, n = 6 X6632; RET: n = 8 DMSO, n = 7 X6632; YUMM 1.7: n = 8 DMSO, n = 8 X6632) *P ≤ 0.05; **P ≤ 0.01; ***P ≤ 0.001.

Finally, we tested the effects of X6632 on tumor-induced angiogenesis following subcutaneous injection of the murine melanoma cell lines B16F10, RET and YUMM 1.7 into syngeneic mice. Daily treatment with DMSO or X6632 (18 mg/kg, ip) was started when the tumor volume reached ∼50 mm^3^ (Figure 5D). Blood vessel density in the tumor and adjacent cutaneous tissue was quantified using the pan-endothelial cell marker CD31 (PECAM-1). The data showed that the vessel density was significantly reduced with X6632 treatment compared to vehicle treatment in all three models (Figure 5E, F). These findings suggest that the anti-angiogenic effects of X6632 may contribute to the pronounced inhibitory effect of the compound on melanoma growth that we have previously reported [24].

### X6632 improves the response to anti-PD-1 blockade to suppress murine melanoma growth

A growing body of clinical evidence indicates that combining immune checkpoint inhibition with anti-angiogenic therapy improves antitumor efficacy compared with immune checkpoint inhibition alone, including better survival outcomes for patients with advanced renal cell carcinoma and hepatocellular carcinoma [36]. As our findings demonstrate that X6632 is an angiogenesis inhibitor, we therefore tested the efficacy of anti-PD-1 treatment in combination with X6632 in inhibiting the growth of B16F10 and YUMM 1.7 murine melanomas.

B16F10 or YUMM 1.7 cells were injected subcutaneously into syngeneic mice, then daily treatment with DMSO or X6632 (18 mg/kg, ip) was started when the tumor volume reached ∼30-50 mm^3^ for B16F10 or YUMM 1.7 tumors. In addition, the mice received IgG or anti-PD-1 antibody injections every third day starting with the first day of treatment with DMSO or X6632 (Figure A, B). The B16F10 tumor model did not respond to anti-PD-1 monotherapy. By contrast, X6632 monotherapy (X6632 + IgG) for 8 days significantly reduced tumor growth, consistent with our previous results [24]. Importantly, the combination of X6632 with PD-1 blockade led to a further reduction in tumor volume (Figure 6A).

**Figure 6:**
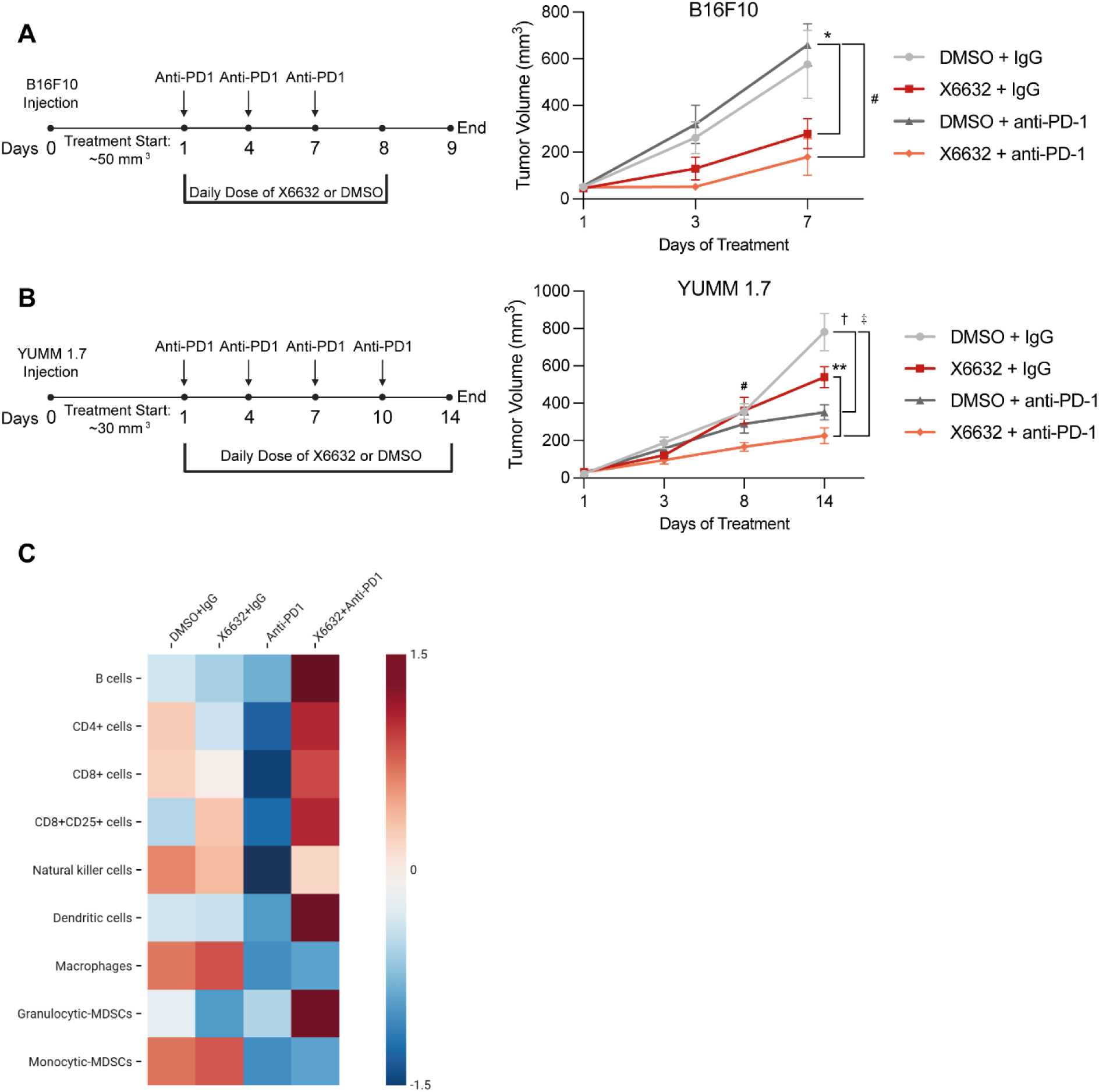
Combination therapy with X6632 and anti-PD-1 blockade in murine melanoma suppression. **A** B16F10 tumor model: tumor cells were injected and daily treatment with DMSO or X6632 (18 mg/kg, ip) was started when the tumor volume reached ∼50 mm^3^. Every third day, mice received anti-PD-1 treatment starting together with DMSO or X6632 treatment. (*, ^#^ P <0.05). **B** YUMM 1.7 tumor model: tumor cells were injected and daily treatment with DMSO or X6632 (18 mg/kg, ip) was started when the tumor volume reached ∼30 mm^3^. Every third day, mice received IgG or anti-PD-1 injections (1 µg/µL, ip) starting together with DMSO or X6632 treatment. (^#, †^, ^‡^, P < 0.05; **P < 0.01). **C** Heatmap displaying the average percentage of immune cell populations identified by flow cytometry in tumors treated with X6632, IgG/anti-PD-1, or their combination, demonstrating that combined therapy preferentially enhances T and B cell infiltration (see Supplementary Fig. 1 for plots of the individual immune cell types).

In the YUMM 1.7 melanoma experiment, tumor-bearing mice received 14 injections of DMSO or X6632, and four IgG or anti-PD-1 treatments (Figure 6B). A significant reduction in tumor growth was observed with anti-PD-1 monotherapy compared to DMSO plus IgG treatment at the end of the experiment, indicating that the YUMM 1.7 model is sensitive to immune checkpoint blockade (Figure 6B). Although X6632 monotherapy reduced tumor growth compared to the DMSO control, this did not reach statistical significance within the timeframe of the experiment. However, a significant reduction in tumor growth was observed when X6632 was combined with anti-PD-1 treatment compared to X6632 monotherapy, and the combination further reduced tumor growth compared to anti-PD-1 monotherapy (Figure 6B), although this did not reach statistical significance within the timeframe of the experiment. Taken together, these results suggest that the combination of X6632 with immune checkpoint blockade elicits enhanced anti-tumor effects compared to monotherapy, and may potentially sensitize unresponsive tumors to immune checkpoint therapy.

To further analyze the immune composition within the treated tumor microenvironment across the different groups, B16F10 melanoma tumors were removed at the endpoint, prepared as single-cell suspensions, and subsequently analyzed by flow cytometry. Tumors treated with the X6632 and anti-PD1 combination exhibited the greatest enrichment of immune cells across all groups (Figure 6C, Supplementary Fig. 1). Specifically, combining X6632 with anti-PD1 drove a pronounced increase in tumor-infiltrating immune cells, including B cells, CD4^+^ T cells, granulocytic-monocyte derived suppressor cells (MDSCs), and activated CD8+ T cells (CD45+CD8+CD25+), which was significantly greater than that observed with anti-PD1 monotherapy (Supplementary Fig. 1). While monocytic MDSCs, dendritic cells, and NK cells did not show any significant changes, the macrophage fraction was reduced in both anti-PD1 and anti-PD1+X6632 treated tumors (Supplementary Fig. 1). Collectively, these findings suggest that X6632 reprograms the immunosuppressive tumor microenvironment of B16F10 melanoma – an immune-cold tumor – toward an immune-permissive phenotype, augmenting anti-PD1 efficacy by promoting CD8+ T cell activation and intratumoral infiltration.

### X6632 inhibits tumor growth and angiogenesis in syngeneic and patient-derived xenograft models of breast cancer

Our previous findings [24] and the results above document the ability of X6632 to suppress melanoma initiation and growth. To extend these findings to other types of cancer, we next tested the effect of X6632 on tumor initiation, tumor growth, and tumor-induced angiogenesis in pre-clinical models of breast cancer. First, we evaluated the effects of X6632 on the survival and proliferation of different human breast cancer cell lines using clonogenic assays. The cells were seeded at a low density and treated with various concentrations of X6632. In all four breast cancer cell lines tested, treatment with X6632 significantly inhibited colony formation compared to the DMSO control, with the most pronounced effect being observed with the SK-BR-3 and BTB474 cell lines (Figure 7A and B).

**Figure 7:**
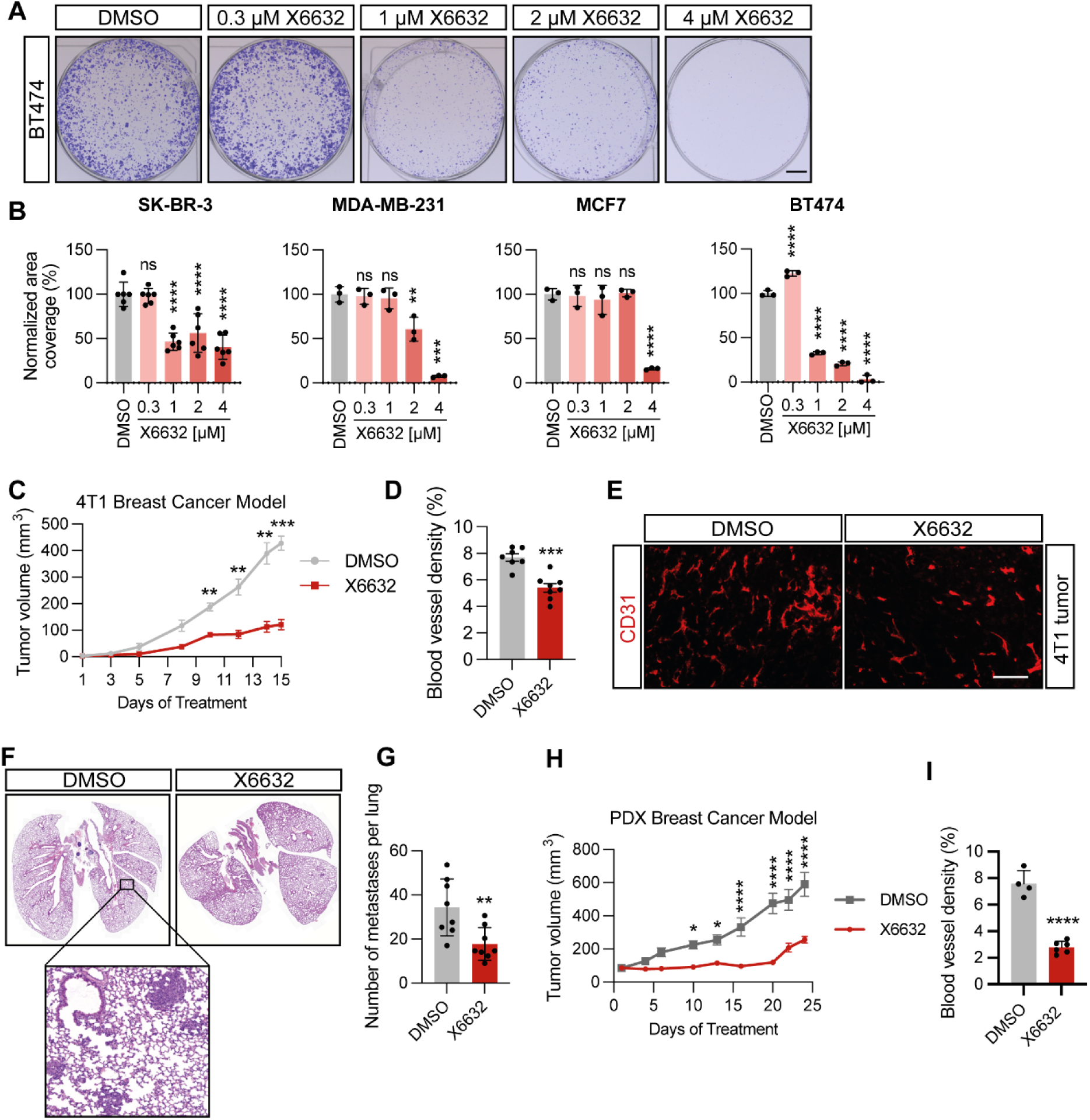
Potent inhibition of growth and angiogenesis in syngeneic and patient-derived xenograft breast cancer models. **A** Representative images of the clonogenic assay using the BTB474 cell line. **B** Different breast cancer cell lines were seeded at a low density and treated with the indicated concentrations of X6632 for 2 to 4 weeks. **C** Mice received daily treatment with DMSO or X6632 (18 mg/kg, ip) for 14 days (n = 8). **D** Quantification of blood vessel density in the tumors after treatment with DMSO or X6632 (n = 8). **E** Representative images of blood vessel density (anti-CD31) in tumor cryosections of animals treated with DMSO or X6632 (scale bar = 100 µm). **F** Representative images of lungs after tail vein injection of 4T1 breast cancer cells and treatment with DMSO or X6632 (18 mg/kg, ip) five times per week for 10 days. **G** Histopathological quantification of the number of metastases per lung in the two different treatment groups. **H** Tumor growth of the HCI-043 PDX model in immune-compromised mice (NOD/*scid*). Mice received daily treatment with DMSO or X6632 (18 mg/kg, ip) for 24 days. **G** Quantification of blood vessel density in the tumors after treatment with DMSO or X6632. X6632 significantly reduced blood vessel density in PDX breast cancer tumors. (DMSO n = 4; X6632 n = 6) ns, not significant (P > 0.05); *P ≤ 0.05; **P ≤ 0.01; ***P ≤ 0.001; ****P ≤ 0.0001.

To explore the impact of X6632 on breast cancer growth in vivo, we employed the 4T1 syngeneic mouse model of triple negative breast cancer (TNBC). First, we examined ID1 and ID3 expression in the 4T1 cells using Western blotting (Supplementary Figure 2), and found that X6632 is able to suppress both baseline and BMP4-induced ID1 and ID3 expression in the cells. Next, we investigated the effect of X6632 on the growth of 4T1 tumors in vivo. Treatment with DMSO or X6632 (18 mg/kg, ip, 14 days) was started when a small tumor was detectable. X6632 treatment significantly reduced tumor growth compared to the DMSO control treatment from 10 days after injection to the endpoint of the experiment (Figure 7C). Furthermore, analysis of blood vessel density within the tumors by immunostaining showed a significant reduction in the X6632 treated tumors compared to the vehicle controls, confirming the findings in the melanoma model (Figure 7D, E). To test the effects of X6632 on metastatic colonization of the lung, 4T1 cancer cells were injected into the tail vein of syngeneic mice, and then treated with DMSO or X6632 (18 mg/kg, ip) five times per week (Figure 7F and G). Histopathological analysis of the lungs revealed that X6632 significantly inhibits metastatic lesion formation compared to the DMSO control.

To extend these findings in a human breast cancer model, we evaluated the anti-tumor efficacy of X6632 in a patient-derived xenograft (PDX) breast cancer model. We utilized the HCI-043 PDX TNBC model which is derived from an untreated primary breast tumor with deletions in BRCA1, BRCA2, and RAD50 [37]. Daily treatment with either DMSO or X6632 (18 mg/kg, ip) for 24 days was started when the tumors reached a volume of approximately 100 mm^3^, and the tumor volume was subsequently measured twice a week (Figure 7H and I). X6632 treatment significantly reduced tumor growth compared to the DMSO control treatment from 10 days after injection to the end of the experiment (Figure 7H). The blood vessel density in the X6632-treated tumors was also significantly reduced compared to the DMSO control (7I). Taken together, these findings demonstrate that X6632 demonstrates potent efficacy in inhibiting breast cancer growth, underscoring its potential for clinical application.

## Discussion

Here we report that X6632 is a potent dual-targeting agent that suppresses tumor growth by acting on both cancer cells and the tumor-associated vasculature. Building on the initial identification of the X6632 compound class as ID1 and ID3 inhibitors [24], our findings demonstrate that X6632 is a coumarin-derived pan-ID protein inhibitor whose activity represents a substantial improvement over the first-in-class ID protein antagonist AGX51 [22–23]. Our data provide broad preclinical evidence in support of potential therapeutic application of X6632 for the treatment of multiple cancer types, as well as for the treatment of diseases involving pathological angiogenesis, such as retinopathies.

ID proteins function as dominant-negative regulators of basic helix-loop-helix transcription factors and ID1 and ID3 have been established as master regulators of cancer cell stemness, tumor aggressiveness, therapy resistance and angiogenesis [2, 14, 18]. AGX51 was the first small molecule ID inhibitor described, which was reported to directly interact with ID proteins, targeting them for proteasomal degradation. It inhibits pathological neovascularization models and exerts anti-tumor activity in breast and colorectal cancer [22–23]. X6632 achieves approximately 10-fold greater inhibitory activity compared to AGX51 in a range of assays, identifying it as a second generation pan-ID inhibitor. AGX51 and X6632 occupy a different chemical space, and in surface plasmon resonance assays we were unable to detect any interaction between X6632 and recombinant ID1 protein (data not shown), suggesting they may suppress ID expression through different mechanisms. Future work will focus on defining the mechanism of action for X6632.

As a pan-ID inhibitor, X6632 can simultaneously target tumor cells and the tumor-associated vasculature. The concept that ID proteins play a decisive role in both tumor cell-intrinsic oncogenic programs and in the regulation of angiogenesis has been appreciated since landmark studies demonstrating that ID1/ID3 double-knockout mice exhibit vascular malformations and fail to support tumor xenograft growth [1, 18]. ID1 and ID3 co-suppression using siRNA in small cell lung cancer cells has been shown to reduce tumorigenicity through both apoptotic and anti-angiogenic pathways [38]. Our data extend these genetic observations pharmacologically, demonstrating that a single agent targeting ID proteins can simultaneously impair cancer cell viability and angiogenic capacity. This dual mechanism is particularly attractive given that resistance to conventional anti-angiogenic therapies frequently arises from compensatory pathways and tumor heterogeneity [39–40].

An important observation from our study is the ability of X6632 to selectively suppress pathological neovascularization without impairing physiological retinal revascularization in the oxygen-induced retinopathy model. The OIR model is the most widely used preclinical model for ischemic retinopathies and serves as a stringent test for anti-angiogenic selectivity, as it simultaneously presents both pathological neovascular tuft formation and physiological vascular regrowth [41]. The finding that a single intravitreal injection of X6632 significantly reduced pathological neovascularization while preserving normal retinal vascular recovery represents a critical therapeutic advantage, consistent with earlier observations made with AGX51 in the same model [22]. This selectivity may reflect the differential expression and functional requirement of ID proteins between pathological and physiological angiogenesis. While ID1 and ID3 are essential for the invasive neovascularization associated with tumors and ischemia, they appear dispensable for the maintenance of normal vascular homeostasis, as evidenced by the minimal expression of ID genes in adult tissues [4, 18]. The therapeutic potential for X6632 therefore extend beyond oncology, and the compound may have utility for the treatment of retinal neovascular diseases such as proliferative diabetic retinopathy, retinopathy of prematurity and the wet form of age-related macular degeneration (AMD).

We have previously shown that X6632 exerts a potent inhibitory effect on the initiation of melanoma in vivo [24]. The ability of X6632 to significantly reduce both the growth of pre-established tumors and tumor blood vessel density in three syngeneic melanoma models further underscores the in vivo anti-tumor and anti-angiogenic activity of this compound. Elevated ID1 expression in melanoma has been associated with decreased patient survival and increased ephrin-A1/EPHA2 signaling [10], and ID proteins play a recognized role in melanoma stemness and BMP-dependent signaling pathways [12, 15]. Beyond melanoma, we demonstrate that X6632 has a broad anti-cancer activity in multiple breast cancer models. X6632 suppressed clonogenic growth in cell lines that represent major molecular breast cancer subtypes, including ER^+^ (MCF7), HER2^+^ (BT474 and SK-BR-3), and triple-negative breast cancers (MDA-MB-231). ID proteins have been implicated in breast cancer growth, invasion, metastasis, stemness and therapy resistance [7–9, 11, 13, 42–43], and our data suggest that pharmacological pan-ID inhibition is effective irrespective of breast cancer molecular subtype. In the 4T1 syngeneic breast cancer model, X6632 treatment reduced primary tumor growth, diminished tumor vasculature, and significantly decreased lung metastasis. The ability of X6632 to potently suppress the growth of two TNBC models in vivo is particularly exciting, given the current difficulties in effectively treating this molecular subtype of cancer [44].

In terms of potential clinical application, the anti-angiogenic properties of X6632 open a number of potential avenues for combination therapies for the treatment of cancer. For example, the combination of angiogenesis suppression with immune checkpoint inhibitors has proven more effective than immune checkpoint inhibitors alone or current standard of care therapies in clinical trials, including for hepatocellular carcinoma, and represents an area of rapidly growing clinical interest [45–46]. The enhanced efficacy observed in the B16F10 and YUMM 1.7 models with the X6632 plus anti-PD-1 combination is consistent with these observations. The abnormal tumor vasculature creates a hypoxic, immunosuppressive microenvironment that hinders T-cell infiltration and favors recruitment of regulatory T cells and myeloid-derived suppressor cells [45, 47]. Judicious anti-angiogenic therapy can transiently normalize tumor vessels, improving perfusion, reducing hypoxia, polarizing tumor-associated macrophages from an immunosuppressive M2-like toward an immunostimulatory M1-like phenotype, and facilitating CD4^+^ and CD8^+^ T-cell infiltration [47–48]. Importantly, lower vascular-normalizing doses of anti-angiogenic agents have been shown to be more effective at enhancing immunotherapy than high doses that cause excessive vessel pruning and exacerbate hypoxia [47]. These results position X6632 as a candidate for combination immunotherapy strategies in melanoma and potentially other tumor types where immune checkpoint inhibitors have shown limited efficacy as monotherapy.

The translational potential of X6632 is further supported by its efficacy in the HCI-043 patient-derived xenograft model, a BRCA1/2/RAD50-deleted TNBC breast cancer model. PDX models are superior to conventional cell line xenografts as preclinical tools in some aspects, as they retain tumor heterogeneity, gene expression profiles, and treatment responses that more closely recapitulate the original patient tumor [49–50]. The significant reduction in tumor growth and vessel density observed over 24 days of treatment in this PDX model suggests that X6632 may be effective against genetically complex, treatment-resistant breast cancers carrying DNA damage repair deficiencies. The demonstration of dual anti-tumor and anti-angiogenic effects in a PDX context strengthens the case for further evaluation of X6632 in clinically relevant translational settings.

From a medicinal chemistry perspective, the coumarin scaffold of X6632 represents a privileged pharmacophore with well-documented anti-cancer and anti-angiogenic properties [51–52]. Coumarin derivatives have been shown to modulate angiogenesis through diverse mechanisms, including suppression of VEGF signaling, inhibition of matrix metalloproteinases, and interference with PI3K/Akt pathways. The identification of X6632 as a specific pan-ID protein inhibitor within this chemical class adds a novel mechanism of action to the coumarin pharmacological repertoire and establishes a structure–activity foundation for further optimization of this compound class toward clinical development.

Ahead of possible clinical application, the pharmacokinetic properties and toxicity profile of X6632 remain to be characterized in detail. With regard to toxicity, no overt signs of toxicity such as piloerection, weight loss, lethargy or abnormal breathing were observed in the animal experiments that we have previously described [24], nor in those reported here, even after more than three weeks of treatment. Furthermore, pathological neovascular tuft formation but not physiological vascular regrowth was inhibited by X6632 in the OIR model. While we demonstrate efficacy across multiple tumor types and models, the long-term durability of responses and potential emergence of resistance mechanisms will require extended treatment studies. The relative contributions of direct anti-tumor versus anti-angiogenic effects in each model remain to be fully dissected, and the optimal dosing schedule for combination with immune checkpoint inhibitors in light of the vascular normalization window concept [47] warrants further investigation.

In conclusion, X6632 is a potent pan-ID protein inhibitor that exhibits dual anti-tumor and anti-angiogenic activity, which represents a significant advance over the first-in-class compound AGX51. Its treatment efficacy in melanoma, breast cancer and ocular neovascularization models, as well as its ability to potentiate the efficacy of immune checkpoint blockade, positions X6632 as a promising candidate for further preclinical development and eventual clinical translation as a multi-modal anti-cancer agent.

## Acknowledgements

We thank Tabea Wagner and Lara Dreyer for excellent technical assistance, as well as Simone Gräßle, Sylvia Vanderheiden and Jana Barylko for the resynthesis of X6632. We gratefully acknowledge the support of the Core Facility Live Cell Imaging Mannheim (LIMa) and the core facility Preclinical Models of the Medical Faculty Mannheim.

## Funding

This work was supported by grants from the Deutsche Forschungsgemeinschaft (DFG, German Research Foundation) to J.P.S. (Project number 259332240/RTG 2099), to J.P.S and Y.S. (Project number 466381716/SL 37/11-1) and from the European Union’s Horizon Europe research and innovation program (grant agreement no. 101096473) under the auspices of the “Lung Cancer-related risk factors and their Impact Assessment” (LUCIA) project. We further acknowledge the scientific data storage service of Heidelberg University (SDS@HD) supported by the Ministry of Science, Research and the Arts Baden-Württemberg (MWK) and the German Research Foundation (DFG) through grant INST 35/1314-1 FUGG and INST 35/1503-1 FUGG.

## Author Contributions

Conceptualization: BKG, JPS. Investigation, Methodology, Formal Analysis: LSG, AD, SS, LD’A, HTW, GS, LPTF, UK, EF, ZR, WT, NJ. Supervision and Resources: ST, SB, CSH, ALW, YS, BKG, JPS. Writing – original draft: LSG, BKG, JPS. Writing – review and editing: AD, WT, CSH, YS, BKG, JPS.

## Data availability

The datasets used or analysed during the current study are available from the corresponding author upon reasonable request.

## Conflicts of Interest

The authors declare no competing interests.

## Notes

### Competing Interest Statement

The authors have declared no competing interest.

