## Supplementary Information for "The coumarin derivative X6632 is a pan-ID protein inhibitor that suppresses tumor growth by targeting cancer cells and the tumor-associated microvasculature"

### Supplementary Figures

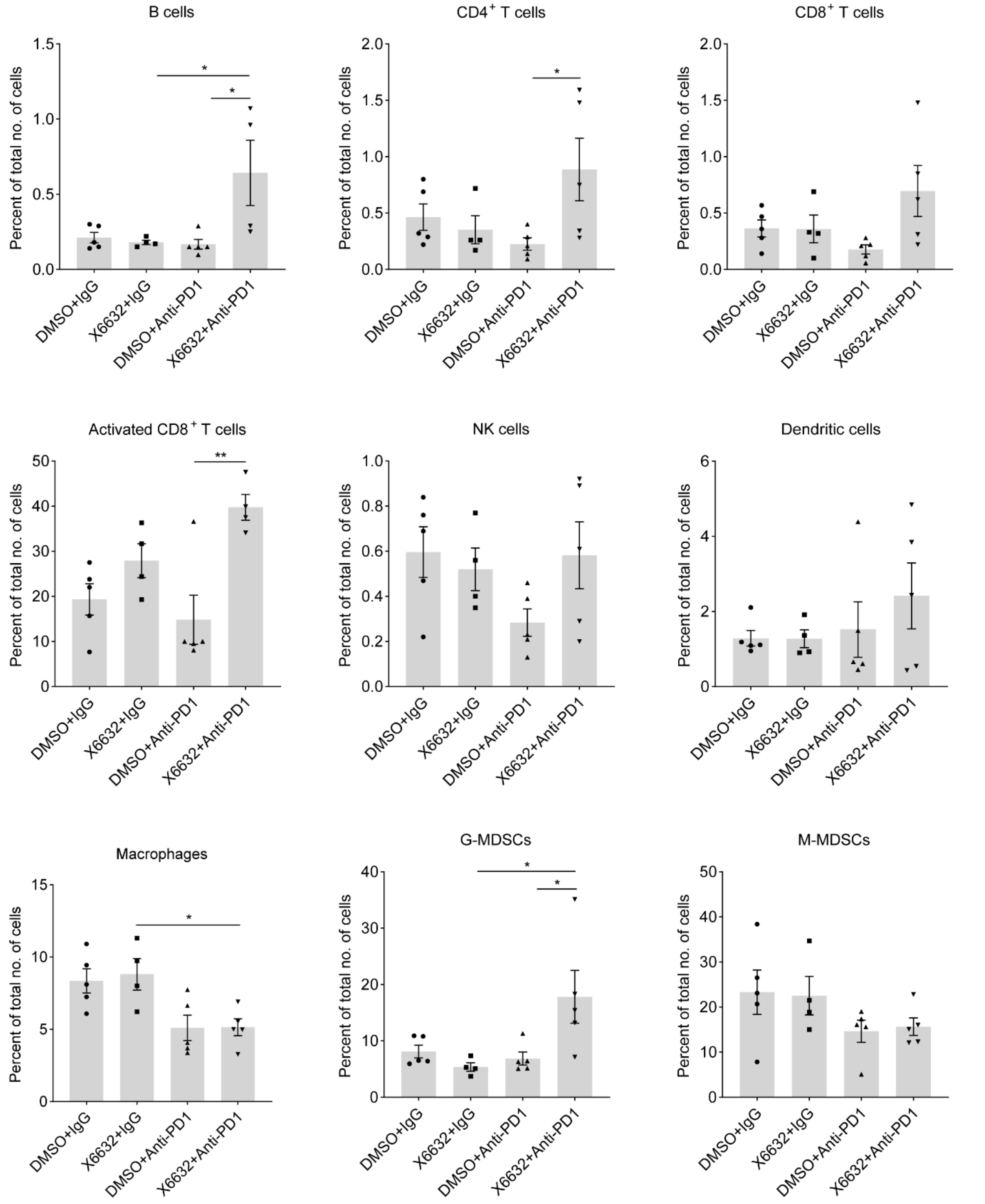


**Supplementary Figure 1. Distribution of immune cell populations identified by flow cytometry in tumors treated with X6632, IgG/anti-PD-1, or their combination.**

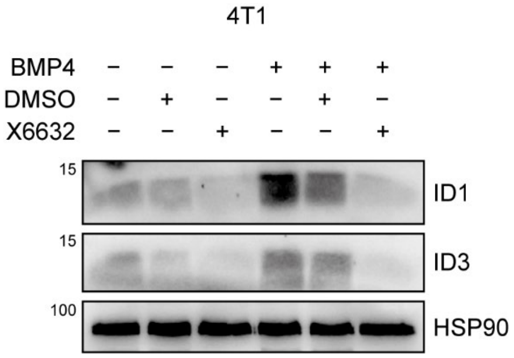


**Supplementary Figure 2. X6632 suppresses BMP4-induced ID1 and ID3 expression in 4T1 breast cancer cells.** 4T1 cells were either stimulated with BMP4 (20 ng/mL) or left unstimulated. In addition, the cells were treated with DMSO (0.1%) or 10 µM X6632 for 24 hours. Evaluation of ID1/ID3 protein expression was carried out using Western blot analysis.

### Supplementary Tables

| Cell types | Surface markers |
|---|---|
| B cells | $CD45^{+}B220^{+}$ |
| CD4+ T cells | $CD45^{+}CD4^{+}$ |
| CD8+ T cells | $CD45^{+}CD8^{+}$ |
| Activated CD8+ T cells | $CD45^{+}CD8^{+}CD25^{+}$ |
| Natural killer cells | CD45+CD49b+ or $CD45^{+}NKp46^{+}$ |
| Dendritic cells | $CD45^{+}CD11c^{+}$ |
| M-MDSCs | $CD45^{+}CD11b^{+}Ly6C^{+}Ly6G^{low}$ |
| G-MDSCs | $CD45^{+}CD11b^{+}Ly6C^{low}Ly6G^{+}$ |
| Macrophages | $CD45^{+}CD11b^{+}F4/80^{+}$ |

**Table S1. Cell surface markers analyzed by flow cytometry to identify immune cell populations.** Single-cell suspensions from tumors were incubated with an antibody mixture and subsequently analyzed by flow cytometry. Cell populations were annotated according to the antibody combinations indicated in the table.

### Supplemetary Methods

#### Synthesis of X6632

The synthesis was carried out according to previously published procedures [1]. Details are available in the Chemotion repository: https://dx.doi.org/10.14272/collection/SGV_2026-07-14 [2].
